# Geometric Protein Optimization

**DOI:** 10.1101/2025.02.10.637504

**Authors:** Dario Wirtz, Claus Horn

## Abstract

The vastness of the space of possible protein variations is often regarded an insurmountable challenge for effective optimization. Traditional approaches focus primarily on substitutions near the binding pocket but are frequently hindered by epistatic effects and non-convex optimization landscapes. Current AI-aided methods can accelerate laboratory procedures but largely target the same substitutions. Here, we propose a complementary approach, Geometric Protein Optimization (GPO), an AI-native framework that fine-tunes the global geometry of the protein by combining a large number of substitutions from diverse locations along the sequence. GPO leverages the strong inverse power-law dependence of electrostatic forces where small adjustments can have large effects on binding affinity. We make the surprising discovery that the inclusion of distal substitutions leads to a smoother and approximately separable optimization landscape. Our empirical investigations reveal three stylized facts about this landscape that we use as a guide to develop BuildUp, a baseline algorithm for GPO. Results show that it is able to navigate this landscape much more effectively and achieve significant improvements in in silico binding affinity (Kd) across diverse protein-ligand complexes. Evaluations of the derived variants through protein-ligand interaction profiling, docking simulations, and molecular dynamics simulations confirm that GPO can achieve beneficial effects even with a sequence-based scoring function.

## Introduction

Efficient protein optimization constitutes one of today’s grand open challenges. Our ability to design proteins with improved fitness has transformative potential for a wide area of applications in biotechnology and medicine, ranging from the enzymatic engineering of efficient biofules to the design of new therapeutic drugs. Traditional protein optimization primarily relies on two approaches[6]: rational design, where domain knowledge is used to target specific residues, often near a binding pocket[14]; and directed evolution, which iteratively applies random substitutions followed by laboratory-based fitness screening[28, 35].

However, both approaches have limitations. Rational design is constrained by its focus on a small set of substitutions, while directed evolution, despite its broader exploration, relies on random substitutions and requires extensive experimental resources. In recent years, semi-rational design, which combines computational predictions with experimental screening, has emerged as a dominant strategy for optimizing protein function[21]. This method leverages structural and functional knowledge to guide mutagenesis, focusing on residues most likely to enhance function, rather than relying purely on random substitutions. Techniques such as site-directed mutagenesis[16], consensus sequence analysis[24], and saturation mutagenesis[25] are widely employed to reduce the search space while maintaining diversity. Advances in machine learning models for protein fitness prediction have further refined semi-rational approaches, enabling more precise identification of beneficial substitutions[36, 5]. While our work differs by focusing on an AI-native optimization framework that explores a broader substitutional space, it aligns with the broader shift toward strategically guided approaches in modern protein engineering.

Based on recent advances in modeling protein structure by artificial intelligence (AI)-based models, like AlphaFold3[12, 1], RoseTTAFold All-Atom[3], and ESM-AA[26, 19, 38], we propose an AI-native framework for in-silico screening, Geometric Protein Optimization (GPO), which significantly enhances the protein optimization process.

For effective in-silico screening, two challenges have to be overcome:

1. Accurately predicting a protein’s fitness for a specific function, and
2. Efficiently searching the vast combinatorial sequence space for optimal designs.

While the first challenge has seen significant progress, achieving near-experimental precision in predicting the binding conformation of protein-ligand complexes[1], and enhanced prediction performance of diverse protein functions[9, 27]. The second challenge remains under-explored and is the focus of our work. Even with an accurate fitness measure, identifying a near optimal protein sequence in the “supra-astronomical” (Francis Arnold, [22]) space of amino acid combinations with 20^*n*^ possibilities, where *n* is the sequence length,^2^ is a daunting, often considered insurmountable task (see e.g. [30]). We hypothesize that an AI-driven optimization approach that combines small effects of a large number of substitutions from diverse locations along the sequence can significantly enhance binding affinity and address the following research question: Given an accurate scoring function, how can state-of-the-art AI models be used more effectively for protein function optimization?

Since an exhaustive search is feasible only for a small number of sites[7], practical approaches often prioritize residues near the binding pocket[15]. However, as our experiments demonstrate, the combination of a large number of substitutions at diverse locations often yields improvements exceeding the effect of binding-pocket substitutions (see Fig. 1), while directed evolution accumulates a large number of suboptimal substitutions. The effects of substituting binding pocket residues in functional wild-type proteins are often negative due to their prior exposure to natural selection. More importantly, their strong effects make the optimization landscape highly non-convex, characterized by epistasis[23].

**Figure 1:**
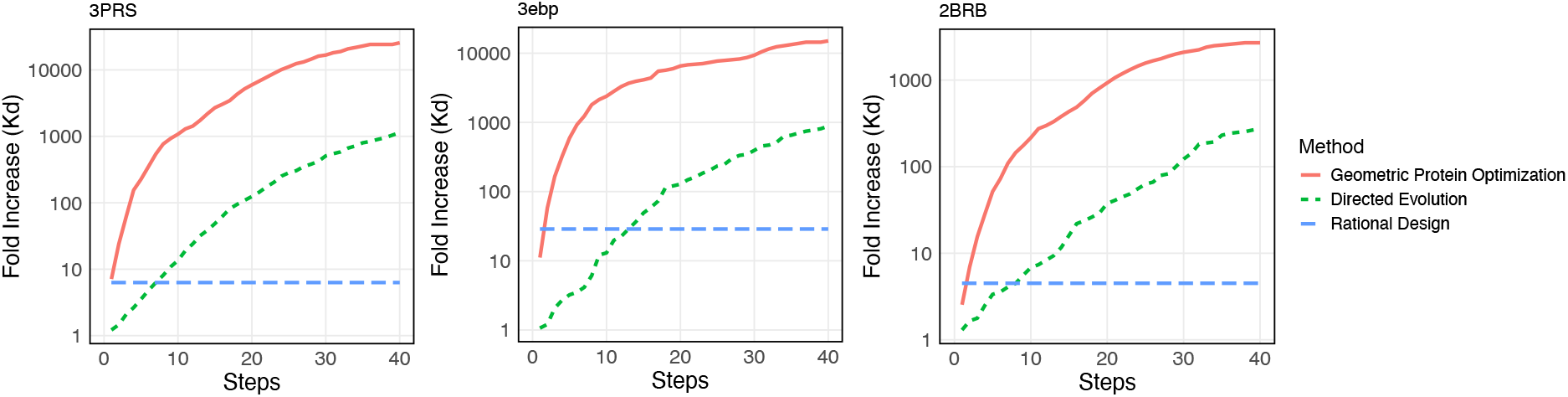
Fold increase in binding affinity achieved by Geometric Protein Optimization (using BuildUp), directed evolution (DE) and rational design (best result obtained from an exhaustive search over the five sites nearest to the ligand) for three different protein-ligand complexes. Both BuildUp and our DE implementation see the same optimization landscape, apply a single substitution per step, and utilize the same number of queries (here 10) to the scoring function per step.

GPO, instead, makes use of state-of-the-art AI-models to combine a large number of substitutions at diverse locations to fine-tune the *global geometry* of the protein. Distal substitutions can have a significant effect on ligand interactions[34, 8, 10] through changes in protein flexibility, structural stabilization, long-range electrostatic effects, or refinements of the binding pocket. The strength of chemical interactions strongly depends on the distance, *d*, between interacting parts[11]. Van der Waals forces decay as 1*/d*^6^, dominating at short distances (3–5 Å), while electrostatic interactions scale as 1*/d*^2^ (monopole-monopole) or 1*/d*^3^ (dipole-dipole), enabling long-range attraction (10–20 Å). Hydrogen bonds, critical for specificity, require precise alignment (2.8–3.5 Å), while charge transfer, important for electron transfer and many enzymatic transformations, even exhibit exponential dependence *e*^−*kd*^.

By analyzing interaction effects and optimization trajectories, we reveal that direct residue substitution measures offer a powerful guide for iterative optimization. Our analysis shows that site and residue effects can be approximately decoupled, reducing the vast protein landscape to a manageable effective size. By prioritizing sites and residues with high direct residue substitution effects, we developed an efficient algorithm that achieves significant improvements in fitness with minimal computational effort.

### Outline and contributions

We measure the effect of single and multiple residue substitutions via a decomposition inspired by Shapley values[29] (details under Methods). Using these measures, we identify three stylized facts for protein optimization that inform the basic design decisions for the BuildUp algorithm presented next. Finally, we describe three experiments using BuildUp that elucidate the effectiveness of GPO.

This paper presents several key contributions:

- We introduce Geometric Protein Optimization (GPO), an AI-native approach that optimizes protein function by leveraging substitutions from diverse locations to fine-tune the global geometry of the protein.
- We identify and validate three key stylized facts of protein optimization that provide new insights into the combinatorial complexity of substitutional landscapes and may guide future improvements in efficient search strategies.
- As a baseline algorithm for GPO, we propose BuildUp, a computationally efficient *discrete gradient* optimization algorithm with favorable scaling behavior, making it ideal for real-world applications in biotechnology and drug design.
- In our experiments, BuildUp achieves significant improvements in in silico binding affinity compared to existing methods.

These findings challenge prevailing assumptions about the intractability of protein optimization and highlight the importance of distal interactions in protein design.

## Results

We utilize the recent BIND model[17] as scoring function, which allows us to estimate the effect of substitutions on binding affinity (more details are given in Methods).

### Stylized Facts of Protein Optimization

We performed experiments by randomly selecting probe sites along the protein sequence and monitor how residue substitution effects at the probe sites change as random substitutions are applied at other sites. We summarize our most important observations about the protein optimization landscape in a list of three stylized facts.

#### SF 1. Residue substitution effects change only gradually

We measure the direct effect of residue substitution by how much the binding affinity (pKd) changes when it is applied. The coefficients of variation (CV = *σ/µ*) are generally around 0.1 (see Fig. 2A). Thus, if residue substitutions at a specific site show potential beneficial effects, this will likely stay so, and the site is unlikely to change its character.

**Figure 2:**
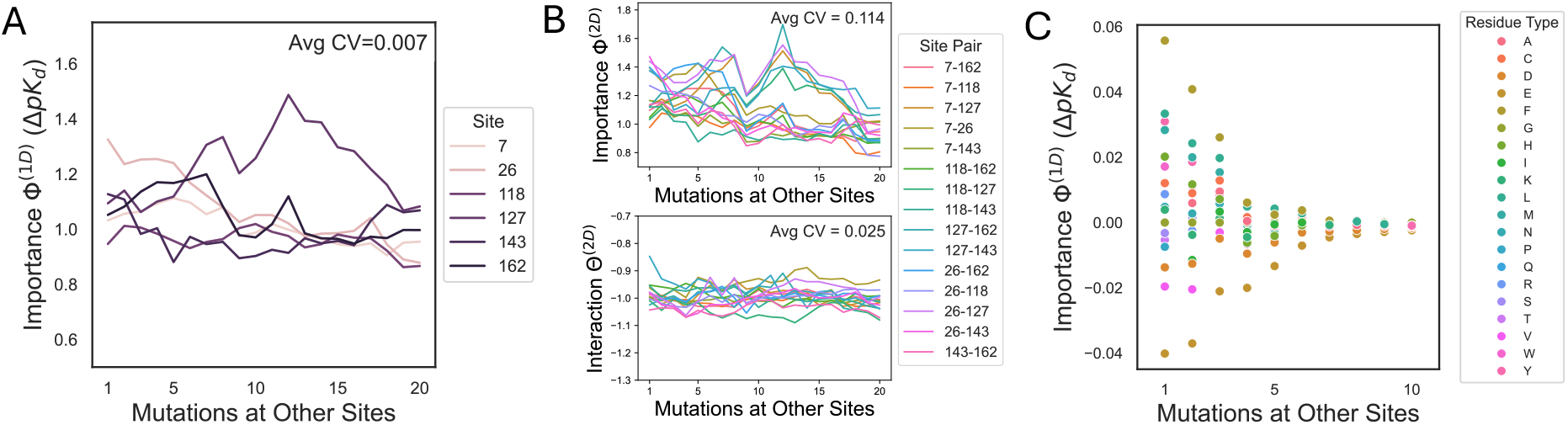
Examples for the stylized facts of protein optimization we observe. **(A)** Evolution of direct residue substitution effects (Φ^(1*D*)^) for randomly chosen sites for the residue with the highest wild-type importance at each site, as substitutions are being applied at other sites. **(B)** Evolution of the effect of pairwise residue substitutions (Φ^(2*D*)^) and pairwise interaction values (Θ^(2*D*)^) for different pairs of residue substitutions as residues are substituted at other sites. **(C)** Change in direct residue substitution effects (Φ^(1*D*)^) for all residues at an unaltered site as residues are substituted at other sites (selected by the BuildUp algorithm).

#### SF 2. Interaction effects are more stable than pairwise substitution effects

We measure the coefficient of variance as above and calculate the average interaction effect over all pairs for a random set of probe sites (see Fig. 2B)). Although the overall effect of pairwise substitutions can vary considerably, the underlying pairwise-interaction values remain more stable, with CV values around 0.02.

#### SF 3. The order of residue substitution effects at a given site stays mostly constant

##### Algorithm 1

BuildUp

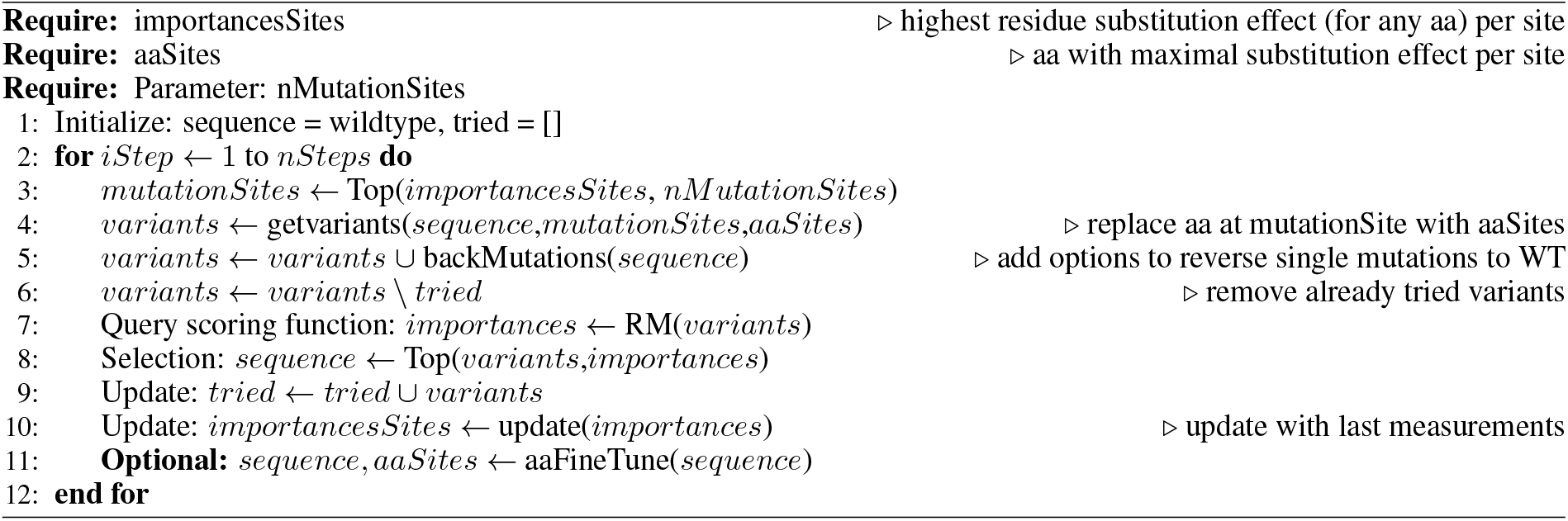

While the mean and variance of the residue-specific substitution effect changes, their ordering only changes slightly (see Fig. 2C)). Thus, to a good approximation, site and residue effects can be factorized and the effect of a residue substitution can be modeled as the product of the site’s position-specific contribution Φ(*s*) and the residue’s effect Φ_*s*_(*a*):

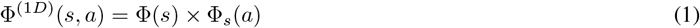

In most cases the dominant effect of a substitution depends on it’s site’s position. We investigate the degree to which this approximation holds below. In combination, these findings (SF1-3) lead us to hypothesize that exhaustive scanning of all possible substitutions at every step is unnecessary and near-optimal substitutions can be efficiently identified by focusing on a small number of site-residue combinations.

### The BuildUp algorithm

BuildUp (see Algorithm 1) is based on the observations reported in the last section. It scans a small set of sites, called *mutationSites*, in each step and selects the one with the highest Φ^(1*D*)^ value (see Methods). The *mutationSites* are chosen as the sites with the highest expected residue substitution effect, as determined by the most recent measured substitution effect at that site. Only the substitution with the highest effect is considered at each site (stored in the variable *aaSites*), with the option of fine-tuning them in a separate subroutine. This approach significantly simplifies the protein optimization problem by reducing the dimensionality of the search space. To continue exploring from local optima and avoid infinite loops, the options for returning to the wild-type residue are included, while variants already on the trajectory are excluded as substitution targets.

#### Performance Comparison

We test BuildUp on a selection of protein-ligand complexes and compare it to directed evolution and AdaLead[31]. Table 1 shows the fold increase achieved by BuildUp in comparison with other algorithms. BuildUp significantly outperforms existing approaches, achieving improvements of more than an order of magnitude in some cases. In addition, it offers fast runtimes and favorable scaling behavior (see Table 2). Since a fixed number of mutationSites is considered at each step, execution time scales linearly with the number of steps and is independent of the length of the protein, enabling fast execution also for large proteins.

**Table 1:** Fold increase achieved by directed evolution, AdaLead and BuildUp after 40 steps. For directed evolution, the standard deviation of five runs is shown in parenthesis.

| PL Complex | BuildUp (mutationSites = 10) | BuildUp (mutationSites = 30) | Directed Evolution | AdaLead |
| --- | --- | --- | --- | --- |
| 2BRB | 1761.7 | 2697.0 | 274.8 (71) | 1467.0 |
| 3PRS | 16707.3 | 25555.0 | 1137.0 (253) | 3528.0 |
| 3ebp | 11174.2 | 15062.0 | 884.0 (111) | 1586.0 |
| 1o0H | 40.8 | 50.3 | 23.3 (3.9) | 33.3 |
| 3dxg | 32.6 | 28.2 | 23.8 (3.8) | 7.1 |
| 1p1q | 296.6 | 378.9 | 144.9 (33.5) | 80.2 |
| 1bcu | 37060.0 | 26500.0 | 11859.0 (11739) | 16050.0 |

**Table 2:** Runtime for 40 steps on a LS40 GPU and scaling behavior, with the number of optimization steps (*n*_steps_), the number of evaluated substitutions (*n*_mutationSites_ = 10), and the number of variants selected per step (*n*_Candidates_)

| Algorithms | Run Time (sec) | Scaling |
| --- | --- | --- |
| Directed evolution, $n_{\text{Candidates}} = 10$ | 2795 | $n_{\text{steps}} \times n_{\text{Candidates}}$ |
| AdaLead, $n_{\text{Candidates}} = 10$ | 3605 | $n_{\text{steps}} \times n_{\text{Candidates}}$ |
| BuildUp, $n_{\text{mutationSites}} = 10$ | 728 | $n_{\text{steps}} \times n_{\text{mutationSites}}$ |
| BuildUp, $n_{\text{mutationSites}} = 30$ | 2266 | $n_{\text{steps}} \times n_{\text{mutationSites}}$ |

#### Effect of residue fine-tuning

We include a subroutine to fine-tune the residues (denoted as *aa*) of a given sequence. It uses the same efficient approach as the main algorithm but is applied to variations in residues instead of sites (see Algorithm 2). The optimization is restricted to sites where BuildUp has applied substitutions and where any residue has a positive substitution effect for the given sequence.

##### Algorithm 2

aaFineTune

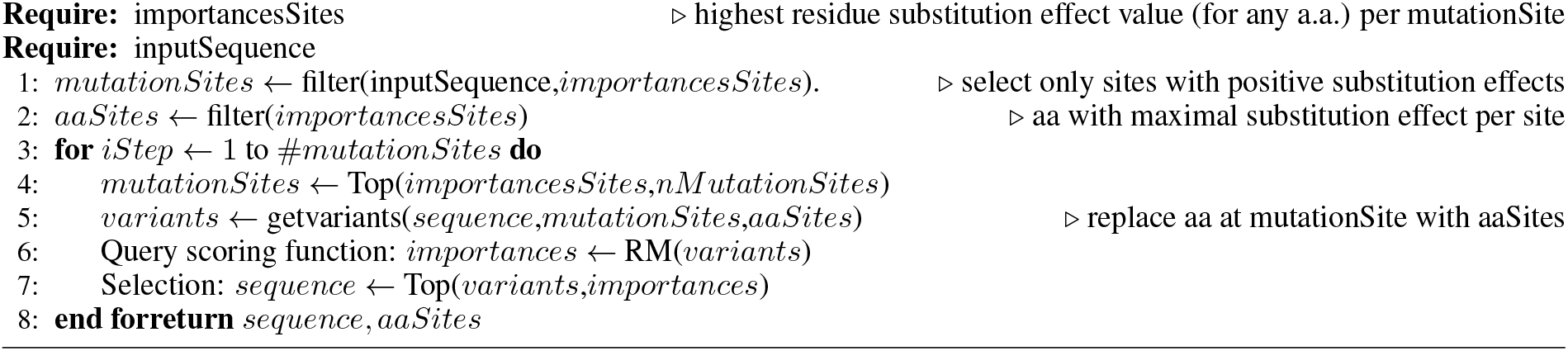

To assess the extent to which SF3 holds, we conduct three measurements. First, we compare the residue substitution effects of other residues not chosen by BuildUp after each sustitution step (Fig. 3A), showing that the wild typ-based residue choice remains a good one, even after a large number of substitutions. These effects could add up if they are not due to uncertainty in the measured values. Thus, in a second experiment, we measure the possible improvement attainable by fin-tuning residues after a larger number of substitutions. We execute the aaFineTune subroutine after 10, 20, and 30 substitutions have been applied by BuildUp, the fraction of residues for which optimization potential exists, i.e. a different residue has a higher score, is only around 5% even after 30 substitutions have been applied (Fig. 3B). The improvement in their substitution effect compared to the amino acid choosen based on wild-type substitution values is usually quite small. The combined effect of optimizing all residuals with *aaFineTune* is similar in size to the average effect of a single substitution step of BuildUp (see Fig. 3C). Thus, although SF3 may not hold universally, particularly for residues with direct interaction at the binding pocket, it effectively reduces the combinatorial complexity of protein optimization without compromising the model’s predictive accuracy in most practical scenarios.

**Figure 3:**
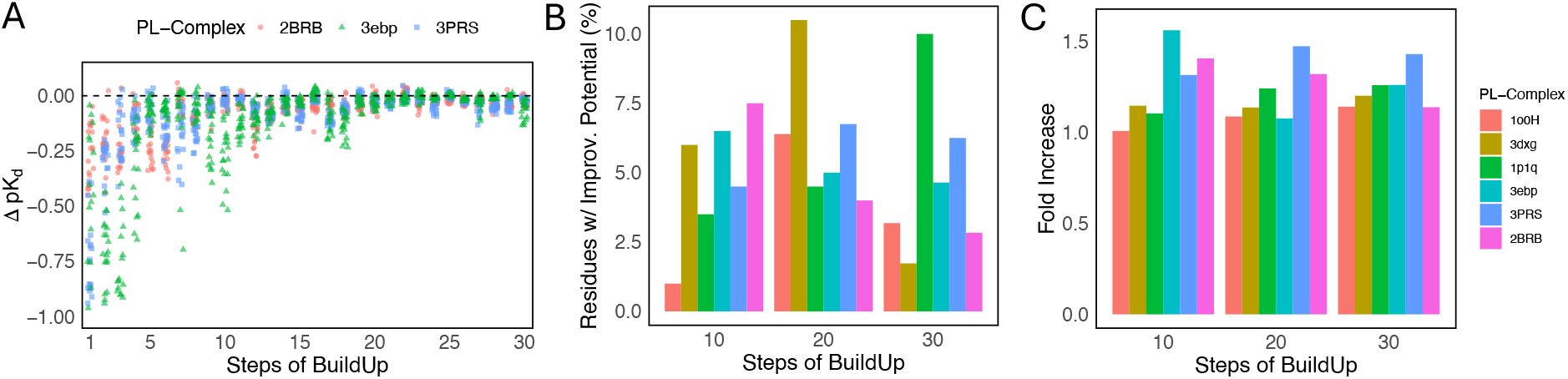
Experimental results confirming the effectiveness of factorizing site and residue optimization: **(A)** Residue substitution effects of other residues compared to the original chosen one at the substitution site selected at each step of buildUp execution. **(B)** Fraction of residues with substitution effects exceeding the one of the initial residues. **(C)** Fold increase achieved by the aaFineTune subroutine.

#### Empirical Validation of BuildUp’s Core Assumptions

Figure 4A) shows the maximal magnitude of direct residue substitution effects Φ^(1*D*)^ per site of the optimized sequence after 73 substitutions compared to the wild type substitution effects values for the same residues. The linear fit shows a statistically highly significant positive relationship between initial residue substitution effects (substitutions applied to the wild type) and final effect (substitutions applied to the optimized sequence), suggesting that a site’s initial residue substitution effect can indeed serve as a useful filter to remove sites of low influence, even after 73 substitutions.

**Figure 4:**
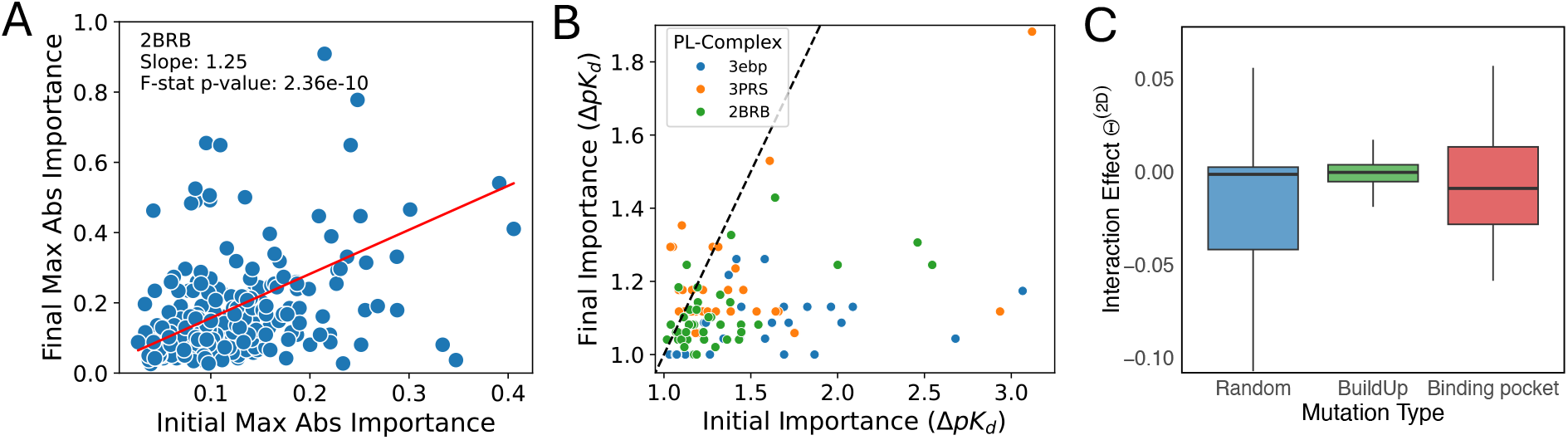
Experiments with BuildUp. **(A)** Site substitution effects, final versus initial, confirming the effectiveness of preselecting sites based on their observed *ϕ*^(1*D*)^ values. **(B)** The large number of substitutions chosen by BuildUp produce *robust* performance improvements through redundancy. **(C)** GPO enables smooth optimization by focusing on substitutions with small interaction effects (Θ^(2*D*)^).

We compare the magnitude of the effect of reverting a substitution from the optimized sequence after 30 BuildUp steps with the magnitude of the initial substitution effect of the same substitution (Fig. 4B). Thus, all substitutions below the identity line have reduced effects. We observe that the advantage of the chosen substitutions is reduced by a large fraction. This shows that BuildUp adds many substitutions that support each other in a partially redundant way, leading to robustly optimized sequences.

### Interactions of residue substitution effects

Our Shapley-inspired decomposition approach (see Methods) allows us to study the relative importance of different substitutional interactions in detail and compare the contributions from direct, pairwise interactions and higher order interactions (Fig. 5), and how they evolve as residue substitutions are being applied by the BuildUp algorithm (see below). We observe that, in most cases, higher order interactions have a destabalizing effect, favoring substitutions that minimize interference and maintain compatibility with one another.

**Figure 5:**
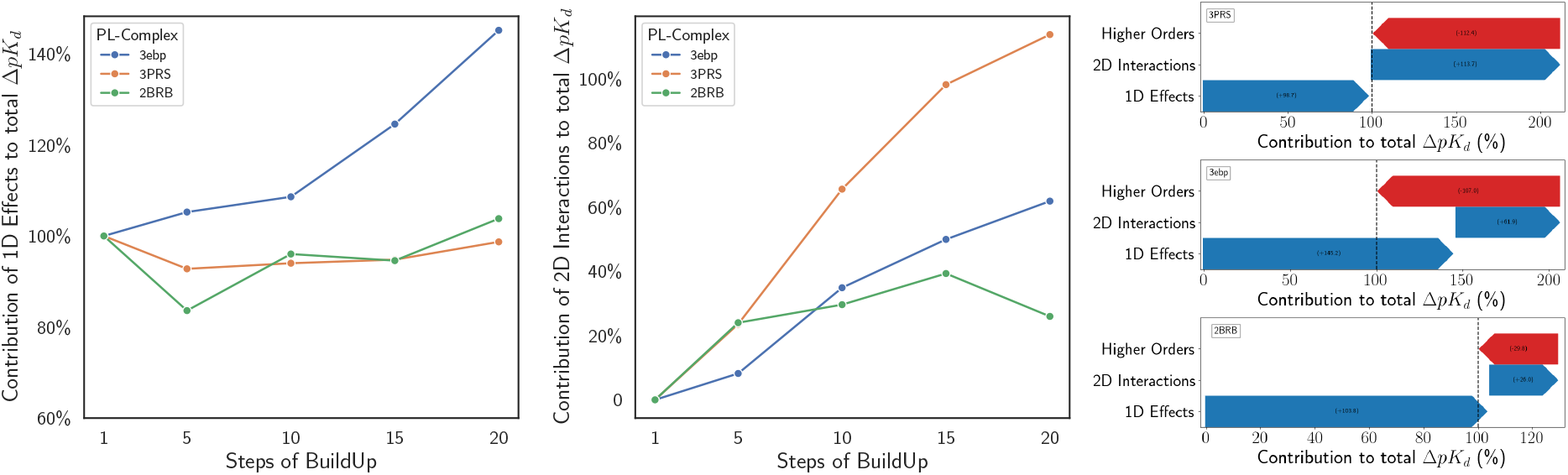
Decomposition of the total Δ*pK*_*d*_ effect enables detailed insight into the relative importance of the interaction of different substitutions and their evolution: **(A)** Percentage of total Δ*pK*_*d*_ that is due to direct effects (Φ^(1*D*)^, as defined under Methods). **(B)** Percentage of total Δ*pK*_*d*_ that is due to pairwise interaction effects (Θ^(2*D*)^). **(C)** Decomposition into direct, pairwise interaction and higher order effects. All for three different protein-ligand complexes.

### Evaluation of optimized protein variants

We evaluate the variants produced by BuildUp through protein-ligand interaction profiling, docking simulations, and molecular dynamics (MD) simulations. Fig. 6A illustrates non-covalent interactions identified by the Protein Ligand Interaction Profiler (PLIP)[2], based on the last frame of 1ns MD simulations. The results show a significant increase in the number of binding connections for the optimized variants. To further validate the optimized variants, we employed AlphaFold3[1] for structure prediction, followed by docking simulations with AutoDock Vina[33]. Results (Fig. 6B) indicate notable improvements, with decreased binding energy values in most cases, reflecting enhanced binding affinity. However, exceptions were observed, such as with 3PRS, likely due to the limited accuracy of docking simulations in capturing complex binding interactions. To investigate this further and refine evaluation, we conducted molecular dynamics simulations (20 replicates with 500’000 steps of 2fs per step = 1ns) and utilized the GEMS model[9] for structure-based protein-ligand affinity prediction. These analyzes revealed that the optimized 3PRS variant exhibits significantly increased binding affinity compared to the wild type (Fig. 6C). Together, these findings suggest that the proposed optimization framework is able to successfully improve binding interactions, notably while only receiving sequence information as input.

**Figure 6:**
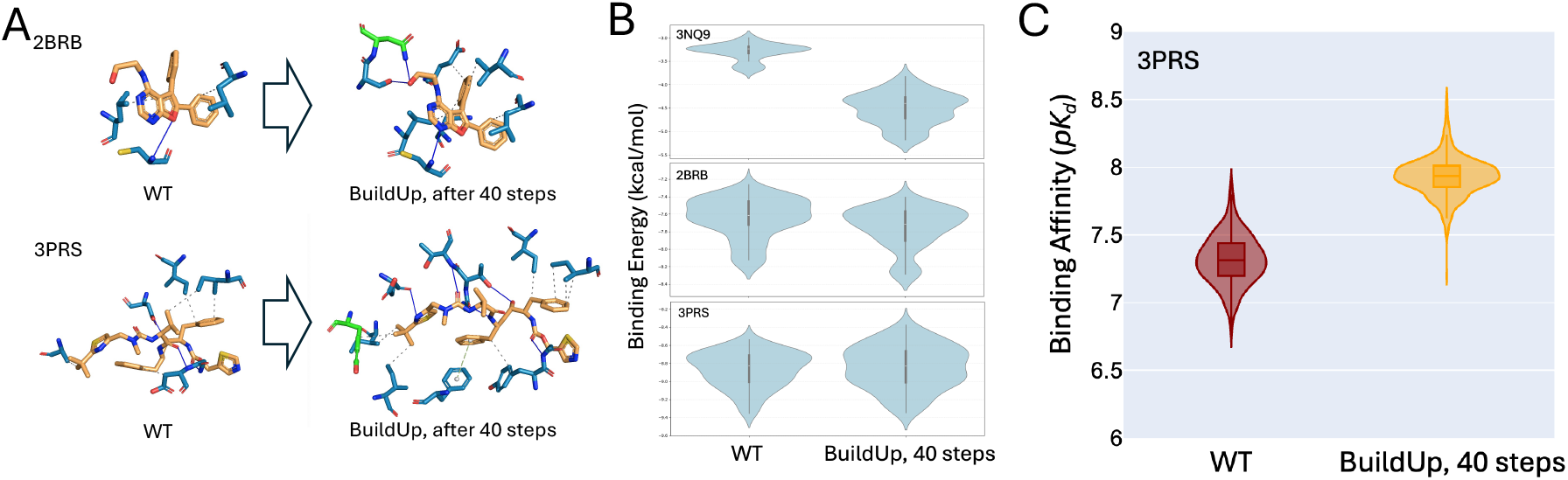
Evaluation of optimized protein variants. **(A)** Increase in interactions identified by PLIP after 40 steps of BuildUp, for 2BRB (top) and 3PRS (bottom). **(B)** Gibbs free energy distributions from AutoDock Vina for 3NQ9 (top), 2BRS (middle), 3PRS (bottom). **(C)** Binding affinity prediction from GEMS for 3PRS.

## Discussion

One can think of BuildUp as implementing a *discrete version of gradient ascent*: it evaluates the local gradient in sequence space by calculating potential improvements along each mutationSite-dimension, selects the direction of the greatest improvement, and iteratively takes steps in that direction. In summary, our observations show that the existence of epistatic effects does not preclude the effectiveness of gradient-based optimization.

To elucidate these findings we propose that the strength of interaction effects may serve as an indicator of the roughness of the optimization landscape. Interestingly, a comparison of the effects of pairwise interactions (Θ^(2*D*)^) between random substitutions, binding pocket substitutions, and those selected by BuildUp reveals significant differences (Fig. 4C). These findings highlight the key to GPO’s success: Selecting substitutions from diverse locations creates a smoother optimization landscape, which enables efficient optimization.

### BuildUp Initialization

The BuildUp algorithm relies on pre-computed residue substitution effect values for all site-residue combinations, which are generated during an initialization step. This initialization process, which takes approximately 1 to 3 hours, is required only once per protein-ligand complex. While this initialization may appear as a significant overhead, it is generally a worthwhile investment, as it enables efficient use of the scoring function, transforming the search process from a random exploration to a guided optimization strategy. Given that practical applications often involve dozens of optimization runs with varying hyperparameters, this upfront computational cost is both reasonable and efficient. Once initialized, subsequent optimization runs are significantly faster, with run-times ranging from 5 minutes to 1 hour for 30 optimization steps.

### Limitations of GPO and BuildUp

In scenarios where substantial improvements can be achieved through multiple binding pocket substitutions, combinatorial optimization methods should be expected to outperform gradient-based algorithms like BildUp. Currently, structure prediction uncertainties are not taken into account during the optimization process. Future iterations of GPO may address these constraints, potentially by integrating experimental feedback. Additionally, hybrid approaches that maintain multiple candidate variants (like directed evolution) could profit from the discovery of high-order interaction effects, at the expense of increased computation.

### Limitations of BIND as a scoring function

BIND is trained on known protein-ligand interactions, which may not generalize well to mutated protein variants and novel or rare binding configurations. Its reliance of ESM2 embeddings instead of explicit modeling of protein structure and ligand docking may limiting its accuracy in scenarios where precise structural interactions are critical. However, our results demonstrate its overall effectiveness in guiding protein optimization.

## Related Work

Existing approaches primarily focus on addressing the combinatorial complexity of binding pocket substitutions, which is complementary to this work. AdaLead[31] is a population-base algorithm for biological sequence design that combines recombination and mutation operations. Its performance compares favorably even to that of much more complex agent-based approaches, making it a competitive baseline for evaluating protein optimization algorithms. In this study, we use AdaLead as the primary reference for comparison. GAMEOPT[4] adopts a game-theoretical approach for combinatorial Bayesian optimization, which focuses on finding equilibria points in the mutational landscape. It factorizes the problem by modeling each site as an independent player. Kirjner et al. introduced a method that derives smoothed fitness landscapes and then learn energy-based and MCMC models in the smoothed landscape[13], which enables protein optimization the limited data regime. Most recently, LatProtRL[18] was introduced, which uses reinforcement learning to optimize protein fitness by traversing a latent space learned using a variant encoder-decoder architecture. Its performance is comparable to AdaLead.

## Conclusions and Future Work

GPO enables a shift from the traditional iterative guess/score/select process to a more *directed* optimization framework, which facilitates more precise, scalable, and data-driven protein engineering. While binding pocket substitutions are often characterized by strong epistatic interactions, giving rise to a combinatorial optimization problem, the small adjustments induced by distal substitutions can be orchestrated efficiently, revealing a potentially large untapped potential for AI-driven protein engineering, complementary to current techniques.

In this work, we demonstrate that, contrary to common belief, *discrete gradient*-based algorithms, like BuildUp, can be implemented efficiently and provide a robust and scalable solution to navigate the landscape of possible protein variations. We anticipate that future improvements in the precision of scoring functions for the predicting of substitution effects will further enhance the practical effectiveness of GPO (e.g. by incorporating structure-based information), and plan to integrate them in future versions. An interesting topic for further investigations will be to disentangle the contributions of distal substitutions in terms of long-range electrostatic interactions, changes in protein flexibility, stabilization of first-shell residues, and structural changes of the binding pocket that facilitate the formation of additional interactions.

Notably, GPO is agnostic to whether input scores are taken from in-silico predictions or laboratory measurements, making it equally applicable to both AI-accelerated design pipelines and experimental-driven optimization workflows. As a result, GPO has the potential to significantly increase the efficiency of protein engineering, by accelerating the discovery of high-functional protein variants and unlocking previously inaccessible regions of the protein landscape. Its insights may be applied to related tasks such as improving enzyme catalysis, optimizing antibody-antigen interactions, enhancing protein stability, and designing novel protein-protein interaction interfaces with a wide number of applications in biochemistry and biomedicine. In future work, we will focus on exploring hybrid approaches that integrate experimental feedback and extending its application to more complex protein engineering challenges.

## Ethical Considerations

Our approach leverages AI-driven optimization to improve protein function, but we recognize the limitations of predictive models in fully capturing the complexity of biochemical interactions. Although computational methods such as GPO provide valuable insights, they should be interpreted as hypothesis-generating tools rather than definitive solutions, and experimental validation remains essential before real-world applications. Furthermore, we acknowledge the potential for dual-use risks in AI-guided protein engineering. To mitigate misuse, we emphasize that our method is designed for beneficial applications, such as enzyme engineering and therapeutic development. We support the implementation of ethical guidelines in AI-driven protein design, ensuring that advances in this field contribute positively to scientific progress and public well-being.

## Methods

The Binding INteraction Determination (BIND) model[17] is a cutting-edge deep learning framework designed to predict protein-ligand binding interactions based on sequence information. Unlike traditional structure-based drug design (SBDD) methods, which require detailed three-dimensional structures for docking and pose prediction, BIND leverages the inherent structure and contextual information encoded in protein language models (pLMs). It was trained on 22,920 protein-ligand interactions from the BindingDB dataset [20]. BIND integrates ESM-2 embeddings with molecular graph representations of ligands using a novel graph-to-transformer cross-attention block, where ligand nodes query protein sequence embeddings to identify potential binding interactions. BIND’s sequence-based approach eliminates the computational overhead of docking, enabling fast exploration of protein space at a fraction of the computational cost of traditional SBDD methods, albeit with comparable prediction performance. In our tests sequence evaluations of took about 0.6 sec on a standard LS40 GPU. The combination of high efficiency with its ability to predict drug-target affinities (*pK*_*d*_ values) with reasonable accuracy makes it ideal for this study of iterative optimization workflows. Ψ defines that fitness, in our case measured by the protein binding affinity *pK*_*d*_, which we use as scoring function.

### Measuring residue substitution effects and interaction effects

Let *M* (*s*, **m**) denote the function that applies a set of substitutions **m** = {*m*_1_, *m*_2_, ߪ}, where each substitution *m* = (*i, aa*) replaces the residue at site *i* of a sequence *s* with the amino acid *aa*. Let Ψ(*s*) denote the fitness function under investigation, and *s*_WT_ the wild type sequence. Inspired by Shapley values[29] we decompose the total effect of a set of substitutions into contributions of single substitutions. The direct effect of a single substitution *m* (1D) is defined as:

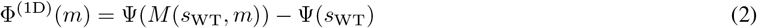

The combined effect of a pair of substitutions *m*_1_ and *m*_2_ (2D) is given by:

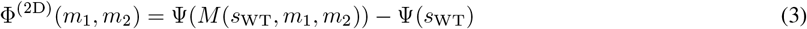

We can then measure the interaction effect between two substitutions, which quantifies their synergistic or antagonistic contribution beyond their individual effects, as:

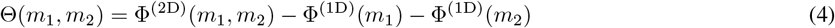

Higher order effects (e.g., Φ^(3D)^, Φ^(4D)^) can be measured analogously by considering the contributions of larger sets of substitutions.

Unlike Shapley values, where effects are averaged over all possible subsets of substitutions, we measure contributions relative to the wild type. This focuses the analysis on sequences near the wild type, improving computational efficiency while retaining the property that contributions sum up to 100%. Specifically, in cases where 3D and higher interactions are absent, for a set **m** of substitutions, the total substitution effect Φ(**m**) satisfies:

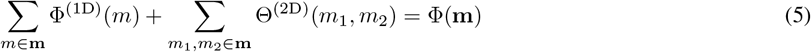

#### Selection of protein-ligand complexes

The protein-ligand complexes used in this study were sampled from the CASF-2016 dataset[32], choosing complexes covering a wide range of affinities (pK 4-8) and containing ligands known to interact with higher affinity with other dissimilar proteins (TM-Score < 0.8)[37]. Complexes without co-factors in the binding pocket and with a binding pocket consisting of a single protein chain were preferred.

## Code availability

The code for this study is freely available on GitHub at https://github.com/claushorn/geometric_protein_optimization.

## Acknowledgements

We thank David Graber and Peter Stockinger for help with evaluations and MD simulations.

## Footnotes

2 For a protein of length 100 the number of configurations exceeds the number of atoms in the universe, for length 200 there are more possibilities than atoms in a universe where every atom is replaced by a full universe of atoms.

